# Alzheimer’s disease risk variant rs11218343 determines functional expression of *SORL1* in microglia

**DOI:** 10.64898/2025.12.15.694337

**Authors:** Malgorzata Gorniak-Walas, Narasimha S Telugu, Ina-Maria Rudolph, Sebastian Diecke, Thomas E Willnow

## Abstract

**Introduction:** Rs11218343 is a non-coding variant of genome-wide significance for sporadic Alzheimer’s disease (AD) with one of the most protective effects known. It localizes to *SORL1*, encoding the AD risk factor SORLA. Still, the functional significance of rs11218343 for AD related processes remains unclear.

**Methods:** We used iPSC lines from donors, or genome-engineered to carry major and minor rs11218343 alleles, to study the impact of rs11218343 genotype on brain cell activities.

**Results:** We show that rs11218343 uniquely controls functional expression of SORLA in microglia, with incrased receptor expression in the minor protective allele correlating with reduced pro-inflammatory responses. This anti-inflammatory effect is seen in donor iPSC lines but not in SNP-engineered isogenic lines, documenting rs11218343 to be diagnostic but not functional.

**Discussion:** Our findings corroborate genetically defined expression levels of *SORL1* in microglia as a determinant of protection from pro-inflammatory stimulation, a function encoded by a haplotype linked to rs11218343.

**RESEARCH IN CONTEXT:** *Systematic review:* Reviewing the literature on single nucleotide polymorphisms (SNP) showing genome-wide association with the risk of sporadic AD, we learned that prior studies identified rs11218343 as a major predictor of protection from the disease. We also learned that this SNP localizes to *SORL1*, encoding the AD risk factor SORLA. However, no data were available whether this SNP controls functional expression of SORLA or other AD-related proteins in brain cell types.

*Interpretation:* Our study documents that rs11218343 controls the expression of *SORL1* in iPSC-derived human microglia, an effect not seen for other microglial genes. Increased expression of SORLA in the minor allele correlates with decreased inflammatory responses in this cell type. These findings suggest that the ability of SORLA to contain pro-inflammatory actions of microglia contributes to its protective effect in AD.

*Future directions:* Our studies document rs11218343 to be diagnostic but not functional in *SORL1* expression control. These findings should be corroborated in a replication cohort of donor cell lines. Also, further analyses need to identify the sequence variation in disequilibrium with rs11218343, that controls *SORL1* gene transcription. Such information will be essential to mechanistically resolve the mode of action of the most protective genotype in sporadic AD known to date.

*HIGHLIGHTS:*

- rs11218343 predicts expressions of AD risk gene *SORL1*
- expression control by the linked haplotype is specific to human microglia
- increased *SORL1* levels with minor allele ameliorates pro-inflammatory responses

## INTRODUCTION

Sortilin-related receptor with A type repeats (SORLA), encoded by *SORL1*, is a type-1 transmembrane receptor expressed in multiple mammalian cell types, including neurons and microglia (reviewed in (1)). It acts as an intracellular sorting receptor that guides trafficking of the amyloid precursor protein (APP) as well as other target proteins between Golgi, cell surface, and endo-lysosomal compartments (2, 3). Previously, *SORL1* gained considerable attention as it shows association with both familial (4-6) but also late-onset sporadic forms of Alzheimer’s disease (AD) (7-10). Recent studies documented aberrant receptor sorting as well as lysosomal dysfunction with coding variants of *SORL1* found in familial AD cases (11-14). While these findings provided important support for the relevance of SORLA-dependent protein sorting in aging brain health, coding variants of *SORL1* as a cause of familial AD are rare. By contrast, non-coding *SORL1* variants of genome-wide significance for risk of sporadic AD are more common, but their functionality remains poorly explained.

Here, we focused on rs11218343, a non-coding variant (T>C) of genome-wide significance for late-onset AD that localizes to intron 21 of *SORL1*. Frequencies of the minor C allele vary between 4% and 30% in European and Asian populations (8, 15). Rs11218343 is unique amongst known single nucleotide polymorphisms (SNP) of genome-wide significance as it shows one of the most protective effects across multiple ethnicities, representing a 19% reduced AD risk for carriers of the C allele (16). Rs11218343 localizes to a distal enhancer region, linked to several microglia gene promoters by proximity ligation-assisted ChIP-seq. Thus, it was predicted to regulate microglial gene signatures that may include *SORL1*, but also other microglia-specific genes (17). However, this hypothesis has not been corroborated experimentally so far.

Using human induced pluripotent stem cell (iPSC)-derived cell models from donors or SNP genome-edited, we now document that a haplotype linked to rs11218343 uniquely determines expression and anti-inflammatory action of SORLA in microglia, arguing for expression in this brain cell type to contribute to genetically determined protective effects of this protein in AD.

## METHODS

### iPSC culture and differentiation

All human induced pluripotent stem cell (iPSC) donor lines used in this study are listed in the Human Pluripotent Stem Cell Registry. Cell lines BIHi005-A (https://hpscreg.eu/cell-line/BIHi005-A), MDCi053-A-49 (https://hpscreg.eu/cell-line/MDCi053-A-49), BIHi242-C (https://hpscreg.eu/cell-line/BIHi242-C), BIHi004-B (https://hpscreg.eu/cell-line/BIHi004-B), BIHi013-A (https://hpscreg.eu/cell-line/BIHi013-A), MDCi240-A (https://hpscreg.eu/cell-line/MDCi240-A), MDCi240-B (https://hpscreg.eu/cell-line/MDCi240-B), and MDCi241-A (https://hpscreg.eu/cell-line/MDCi241-A) were provided by the Stem Cell Facility of the MDC, Berlin. Line BIHi043-A was generously provided by the Institute of Diabetes and Regeneration Research, Munich (https://hpscreg.eu/cell-line/HMGUi001-A). Line TMOi001-A is commercially available (Thermo Fisher Scientific, https://hpscreg.eu/cell-line/TMOi001-A). All cell lines were subjected to initial quality control, including SNP karyotyping (18), and routinely tested to be free of mycoplasma contamination. Generation of isogenic *SORL1* rs11218343 T>C iPSC line by CRISPR/Cas9-based editing as well as *SORL1* and *APOE* genotyping protocols are detailed in the supplementary method section.

Culture and differentiation of iPSC lines into neurons (19), astrocytes (20), or microglia (21) were performed according to published protocols, with adaptations described in the supplementary methods. Protocols for gene and protein expression analysis using quantitative RT-PCR and Western blotting are also given in the supplementary methods.

### Statistical analysis

Data are presented as mean ± standard deviation (SD). Statistical significance of data was determined by nested *t* test when comparing genotype groups, or Student’s *t* test when comparing individual T/C or C/C cell lines to a reference T/T line as stated in the respective figure legends.

## RESULTS

To interrogate the impact of rs11218343 on *SORL1* expression (Fig. 1A), we studied human induced pluripotent stem cell (iPSC) lines from donors with T/T (4 lines) or T/C (5 lines) genotype (Fig. S1A-B). All donor cell lines were homozygous for *APOEε3* (Fig. S1C). SNP genotype did not impact expression of pluripotency markers (Fig. S1D). It also did not affect the robust levels of *SORL1* transcript (Fig. 1B) or SORLA protein (Fig. 1C-D) seen in stem cells.

**Figure 1.**
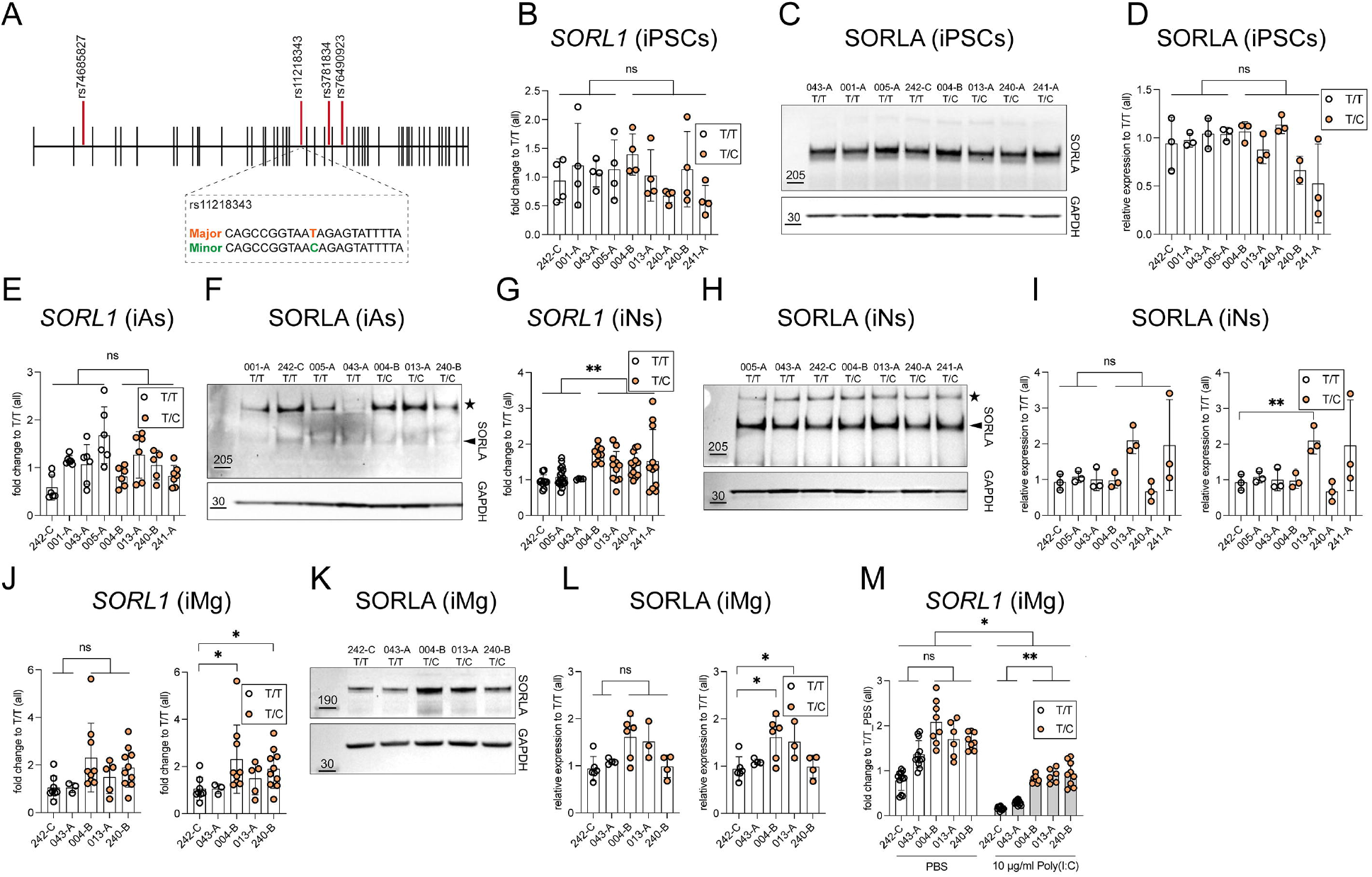
Rs11218343 determines *SORL1* expression in neurons and microglia from donor iPSC lines. **(A)** SNPs in human *SORL1* associated with sporadic AD at genome-wide significance (from alzforum.org/mutations/sorl1-haplotype). Sequence variants for rs11218343 are given below. **(B)** Quantitative RT-PCR analysis of *SORL1* transcript levels in donor iPSC lines. Data are as mean ± SD of 4 biological replicates (from 4 independent differentiations) for each donor line. Statistical significance of data was determined using nested *t* test comparing genotypes. ns, not significant (**C, D**) Levels of SORLA in lysates of iPSC lines of the indicated genotypes were quantified by Western blot analyses (C) and densitometric scanning of replicate blots (D). Relative levels of SORLA in each sample was normalized to the GAPDH loading control in each sample. Data are as mean ± SD of 2-3 biological replicates (from 2-3 independent differentiations) for each donor line. Statistical significance of data was determined using nested *t* test comparing genotypes. (**E**) *SORL1* transcripts in induced astrocytes (iAs) carrying T/T or T/C genotypes of rs11218343. Data are mean ± SD (5-7 biological replicates), expressed as fold change compared to the mean of T/T lines (set to 1, nested *t* test). (**F**) SORLA levels in iAs lysates. The star marks a non-specific band. GAPDH served as loading control. (**G**) *SORL1* transcripts in induced neurons (iNs) carrying T/T or T/C genotypes. Analysis performed as in (B) from 4-18 biological replicates. **, p < 0.01. (**H, I**) SORLA levels in iNs determined by Western blot (H) and densitometric scanning of replicate blots (I). Data are mean ± SD (3 biological replicates). Significance determined by nested *t* test (comparing genotypes, left) or Student’s *t* test (comparing all lines to line 242-C, right). (**J**) *SORL1* transcript levels in induced microglia (iMg) carrying T/T and T/C genotypes. Data analysis as in (B) from 3-10 biological replicates. Significance determined as in (I). *, p < 0.05. (**K, L**) SORLA levels in iMg determined as by Western blot (K) and densitometric scanning of replicate blots (L). Data are mean ± SD (4-7 replicates). Significance determined by nested *t* test (left) or Student’s *t* test comparing individual lines to line 242-C (right). (**M**) *SORL1* transcript levels in iMg treated with PBS or 10 μg/ml poly(I:C). Analysis as in (B) from 6-12 biological replicates per cell line.

Next, we differentiated donor iPSC lines into induced astrocytes (iAs, Fig. S2A-B), neurons (iNs, Fig. S2D-E), or microglia (iMg, Fig. S2G-H), resulting in the expected decrease in pluripotency markers and a concomitant induction of astrocyte (Fig. S2C), neuron (Fig. S2F), or microglia (Fig. S2I-J) specific genes. Comparable levels of *SORL1* transcript were detected in iAs of both genotypes (Fig. 1E) but did not translate into appreciable amounts of SORLA protein (Fig. 1F). By contrast, levels of *SORL1* transcripts were increased in iNs with T/C as compared to T/T genotypes (Fig. 1G). Increased levels of transcripts translated into higher levels of SORLA protein in 2 of the 4 T/C iNs lines (Fig. 1H-I), yet overall protein levels remained low in both genotypes. Levels of *SORL1* transcript (Fig. 1J) were also increased in 2 of 3 T/C lines when differentiated to iMg and translated in robustly increased levels of SORLA protein in these lines (Fig. 1K-L). An association between rs11218343 genotype and microglial expression levels of *SORL1* was further confirmed in iMg treated with poly(I:C), a pro-inflammatory stimulus known to reduce *SORL1* transcript levels (22). *SORL1* transcript levels were always higher in T/C compared to T/T genotypes, both under basal and under poly(I:C)-treated conditions (Fig. 1M).

To test functional causality for rs11218343, we used genome editing to convert T/T donor lines 005-A (Fig. S3A-B) and 053-A (Fig. S4A-B) to isogenic lines homozygous for C/C. No impact of genotype conversion on expression of pluripotency markers (Figs. S3C and S4C), or levels of *SORL1* transcript (Figs. S3C and S4C) or protein (Figs. S3D-E and Fig. S4D-E) were seen. Subsequently, isogenic clones of 005-A were differentiated into iNs, showing comparable appearance (Fig. S3F) and induction of neuronal gene expression (Fig. S3G) in T/T and C/C lines. As donor line 005-A failed to generate iMg, the impact of rs11218343 on microglial gene expression was tested in isogenic clones of donor cell line 053-A, showing comparable microglial morphology (Fig. S4F) and marker expression (Fig. 4G-H) in T/T and C/C variants. Contrary to the findings in donor cell lines, neither levels of *SORL1* transcript (Fig. 2A, D) nor SORLA protein (Fig. 2B-C, E-F) were impacted in iNs or iMg by the presence of the C allele when comparing isogenic T/T and C/C genotypes. These findings argued rs11218343 to be predictive but not functional of *SORL1* expression control. The activity of exemplary microglial gene promoters, spatially linked to the enhanced region harboring rs11218343 (17), showed comparable transcript levels in iMg generated from donor (Fig. 2G) or isogenic cell lines (Fig. 2H) of T/T, T/C, and C/C genotypes, arguing that this SNP does not predict global microglial gene signatures but shows specificity for *SORL1*.

**Figure 2.**
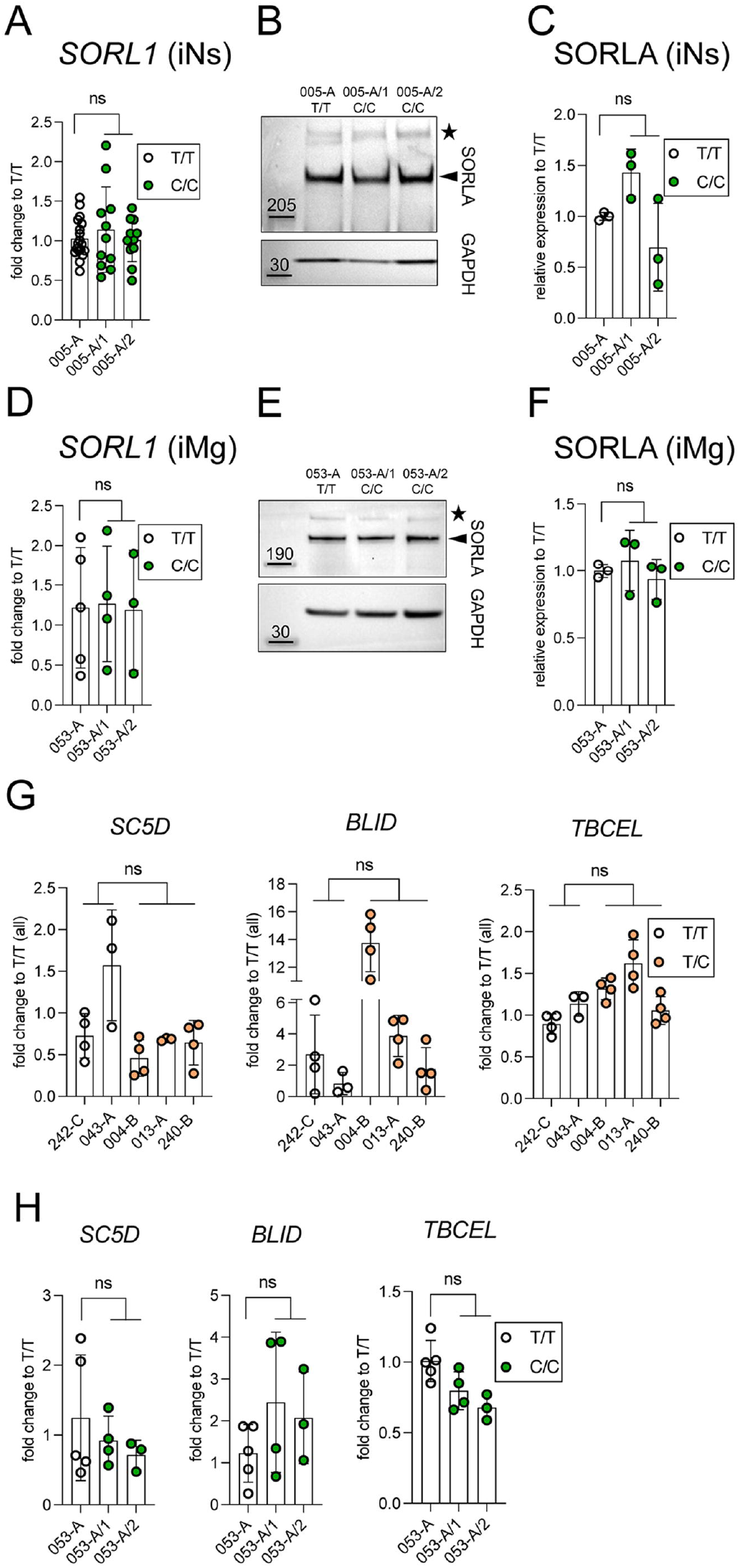
Rs11218343 does not predict *SORL1* expression levels in neurons and microglia from genome-edited isogenic cell lines. (**A**) Quantitative RT-PCR analysis of *SORL1* transcript levels in induced neurons (iNs) derived from parental iPSC line 005-A (T/T) and two independent isogenic cell clones, genome-edited to carry the C/C genotype (005-A/1, 005-A/2) on day 14 of culture. For each cell line, data are given as mean ± SD of 11-18 biological replicates (from 8-11 independent differentiations), expressed as fold change compared to the T/T cell line (mean set to 1). Statistical significance of data was determined using nested *t* test comparing the two genotype groups. ns, not significant. (**B**) Western blot analysis of SORLA levels (arrowhead) in lysates of iNs of the indicated genotypes. The star marks a non-specific band reacting with the SORLA antiserum. Detection of GAPDH served as loading control. (**C**) Levels of SORLA in iNs lysates were determined by densitometric scanning of replicate Western blots (as exemplified in B). Relative expression of SORLA was normalized to GAPDH in the respective sample. Data are given as mean ± SD of 3 biological replicates (from 3 independent differentiations) for each cell line. Significance of data was determined using nested *t* test comparing the two genotype groups. (**D**) Quantitative RT-PCR analysis of *SORL1* transcript levels in iMg derived from parental iPSC line 053-A (T/T) and two independent isogenic cell clones, genome-edited to carry the C/C genotype (053-A/1, 053-A/2) on day 38 of culture. Analysis was performed as in (A) using data of 3-5 biological replicates from 3-5 independent differentiations, expressed as fold change compared to the T/T cell line (mean set to 1). Statistical significance of data was determined using nested *t* test comparing the two genotype groups. (**E, F**) Levels of SORLA in lysates of isogenic iMg lines of the indicated genotypes were determined by Western blot analysis (E) and densitometric scanning of replicate blots (F). Relative expression levels of SORLA were normalized to the GAPDH loading control in each sample. Data are shown as mean ± SD of 3 biological replicates (from 3 independent differentiations) for each cell line. Statistical significance of data was determined using nested *t* test comparing the two genotypes. (**G**) Quantitative RT-PCR analysis of transcript levels for *SC5D, BLID*, and *TBCEL* in induced human microglia (iMg) derived from donor iPSC lines of the indicated T/T and the T/C genotypes (day 38 of culture). For each donor cell line, data are given as mean ± SD of 3-4 biological replicates (from 3-4 independent differentiations), expressed as fold change compared to all T/T cell lines (mean set to 1). Statistical significance of data was determined using nested *t* test comparing the two genotypes. ns, not significant. (**H**) Analysis as in panel (G) comparing transcript levels of *SC5D, BLID*, and *TBCEL* in iMg derived from isogenic iPSC lines of the indicated T/T and the C/C genotypes (day 38 of culture). Significance of data was tested by nested *t* test comparing the two genotypes (mean ± SD of 3-5 biological replicates from 3-5 differentiation experiments) for each donor line.

Given the more robust receptor levels in iMg as compared to iNs, we focused on microglia to interrogate the predictive value of rs11218343, not only for expression but also for activity of SORLA. Here, we exposed iMg derived from donor cell lines to pro-inflammatory stimuli by poly(I:C), an activity related to the ability of SORLA to control functional expression of the pattern recognition receptor CD14 (22). Microglia with T/C genotype showed a decreased pro-inflammatory response as documented by reduced levels of cytokines released following poly(I:C) stimulation. This impacted response was most pronounced in 2 of the 3 T/C donor lines (Fig. 3A). A reduced pro-inflammatory response was due to reduced levels of cytokine gene transcription (Fig. 3B), in line with the mode of poly(I:C) action on pro-inflammatory gene expression (23). Consistent with comparable *SORL1* expression levels, pro-inflammatory responses to poly(I:C) were similar in isogenic T/T and C/C lines (Fig. 4A-B).

**Figure 3.**
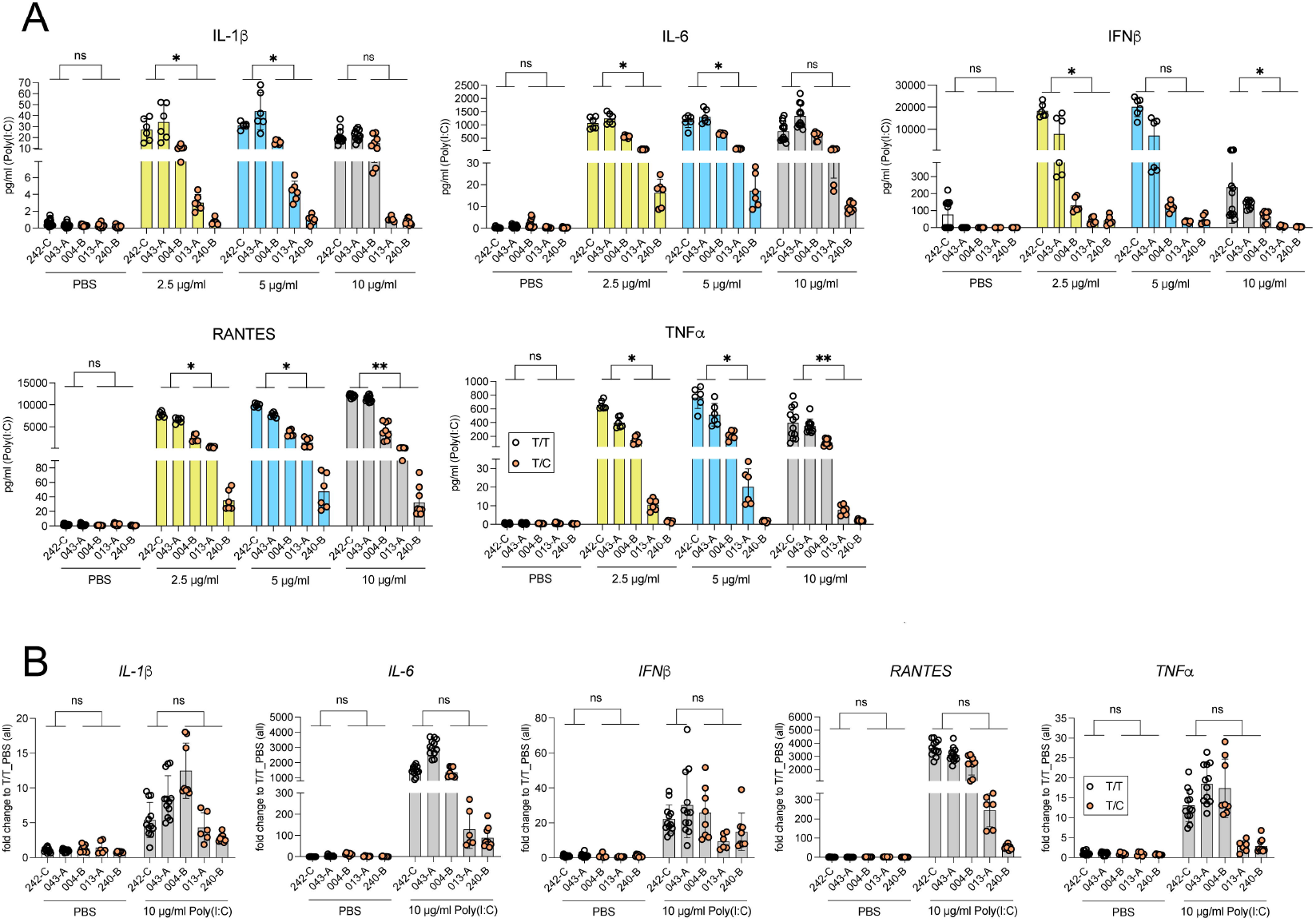
Rs11218343 predicts pro-inflammatory cytokine release by induced microglia from donor cell lines. (**A**) Cytokine levels in supernatants of iMg lines carrying T/T or T/C genotypes treated with PBS or poly(I:C). Data are mean ± SD (6-18 biological replicates, nested *t* test). (**B**) Cytokine transcript levels in iMg carrying T/T or T/C genotypes treated with PBS or 10 µg/ml poly(I:C). Data are mean ± SD of 6-12 biological replicates (nested *t* test). ns, not significant; *, p < 0.05; **, p < 0.01.

**Figure 4.**
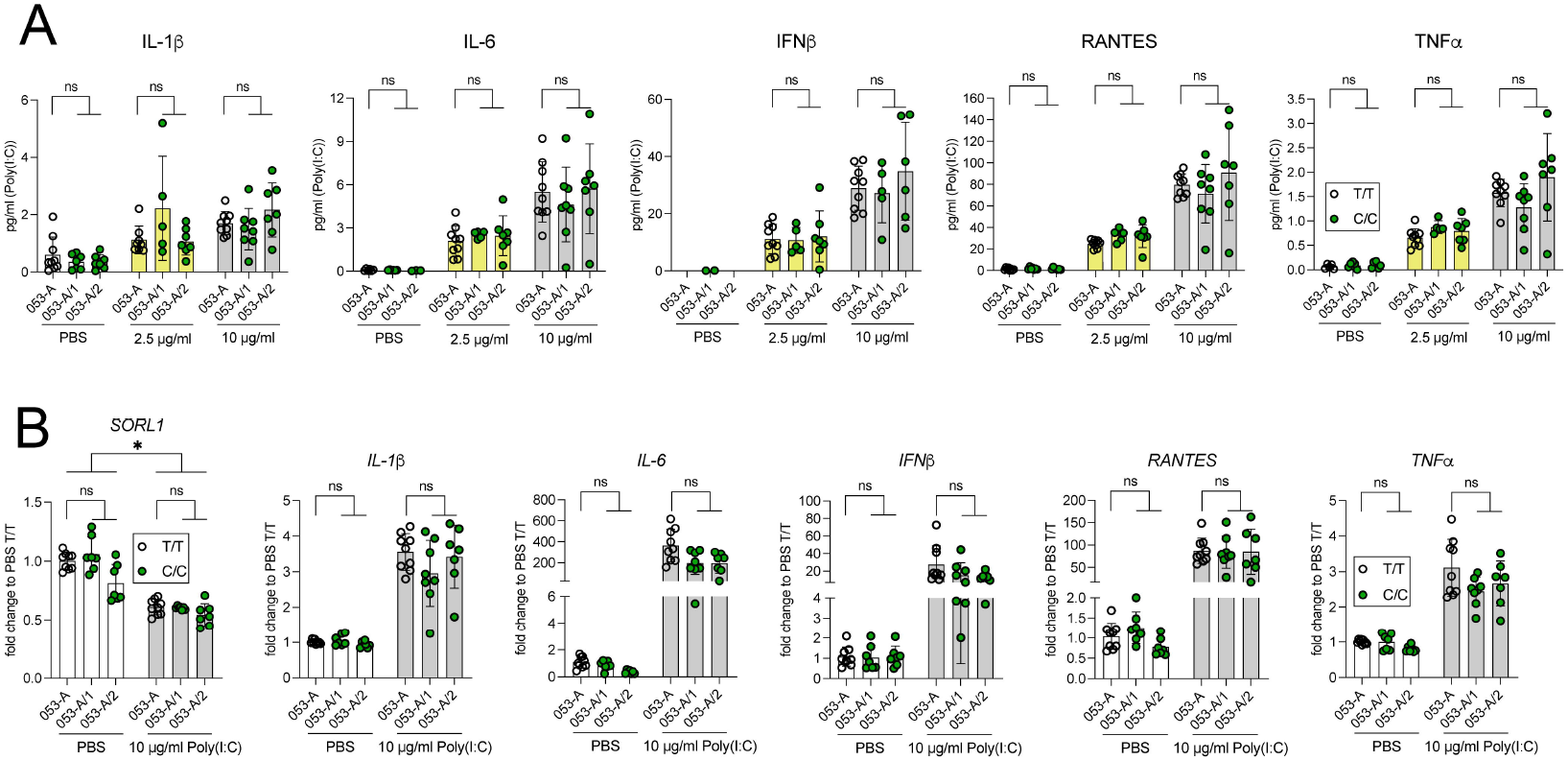
Rs11218343 does not predict pro-inflammatory responses of induced microglia from genome-edited isogenic iPSC lines. (**A**) Cytokine levels in supernatants of iMg derived from isogenic iPSC lines of the indicated T/T and C/C genotypes treated with PBS or poly(I:C). Data are mean ± SD (5-9 biological replicates, nested *t* test). (**B**) Cytokine and *SORL1* transcript levels in iMg derived from isogenic iPSC lines of the indicated T/T and C/C genotypes treated with PBS or 10 µg/ml poly(I:C). Data are mean ± SD of 7-9 biological replicates (nested *t* test). ns, not significant; *, p < 0.05.

## DISCUSSION

According to current hypotheses, intracellular sorting of cargo by SORLA supports several cellular pathways protective against neurodegenerative processes. With relevance to our study, SORLA has been shown to modulate endo- and exocytic activities in human microglia (24, 25), providing arguments for increased receptor expression as determinant of protection against sporadic AD. Early studies provided support for this hypothesis by documenting some *SORL1* risk genotypes, identified in smaller cohorts, to affect translation efficiency (26) or inducibility of this gene by brain-derived neurotrophic factor (27, 28).

Now, our study extends these findings to a *SORL1* genotype of genome-wide significance, associated with risk of sporadic AD. In line with a protective role for SORLA in the disease, the minor allele variant of rs11218343 is associated with increased levels of expression and activity of the receptor. Interestingly, this effect is not seen uniformly in all cell types tested herein. Rather, the predictive value of rs11218343 for SORLA actions seems to be most prominent in microglia, a cell type implicated in neuroprotective actions of this receptor in prior work (22, 24, 25). Interestingly, gene expression control by rs11218343 is not seen for other microglial genes controlled by a microglial enhancer linked to rs11218343, as predicted by *in silico* analyses (17). Because functionality of rs11218343 is seen in donor cell lines but not iPSC lines genome-edited for this SNP, *SORL1* expression control must be exerted through a yet unidentified sequence variation in linkage disequilibrium, or through the entire haplotype. Of note, rs11218343 also seems to predict *SORL1* transcript levels in induced neurons (Fig. 1G). Whether increased gene transcription translates into higher neuronal acivity levels of the receptor is more difficult to assess as, in our hands, SORLA protein levels are typically low in induced cortical neurons as compared to iMg (Fig. 1).

## Supporting information

Sumplementary methods and figures 1 to 4

## Acknowledgement

We are indebted to N. Hübner (MDC) and Fadumo Abdullahi Mohamed (AU) for helpful discussions, and K. Kampf, T. Pasternack, C. Kruse, K.-M. Pedersen, and A. Højland for expert technical assistance. Studies were funded in part by a grant from the Novo Nordisk Foundation (NNF18OC0033928) to TEW.

## Declaration of interest

The authors declare no competing interests.

## Author contributions

MGW, NST, and IMR designed and conducted the experiments and analyzed data. MGW, SD, and TEW conceptualized the study and evaluated data. MGW and TEW wrote the manuscript.

## Notes

### Competing Interest Statement

The authors have declared no competing interest.

