## Supplementary material for "Alzheimer’s disease risk variant rs11218343 determines functional expression of *SORL1* in microglia": Sumplementary methods and figures 1 to 4

**SUPPLEMENTARY METHODS**

**Generation of isogenic *SORL1* rs11218343 T>C iPSC line**

To generate isogenic cell lines homozygous for the minor (C/C) variant of rs11218343, we performed genome editing in induced pluripotent stem cell (iPSC) lines BIHi005-A and in a subclone of BIHi043-4 (MDC i053-A-49), both homozygous for the T/T variant. CRISPR/Cas9-based editing was carried out following previously established protocols (1). In brief, two guide RNAs (gRNAs) flanking the single nucleotide polymorphism (SNP) target site were designed using the Benchling platform and synthesized by Integrated DNA Technologies (IDT). The sequences of the gRNAs were 5′-TAAAATACTCTATTACCGGC-3′ and 5′-CTATTACCGGCTGGGTGCAG-3′. For homology-directed repair, single-stranded oligodeoxynucleotide (ssODN) templates were designed to incorporate the T>C base change at the desired locus. These donor templates were ordered as AltR-HDR single stranded DNA donor templates from IDT, with the following sequences: 5′
GATCCTCCCACCTCGGCCTCCCAAACTGCTGGGATTACAGATGTGAGCCACTGCA
CCCAGCCGGTAACAGAGTATTTTAAAATAACATCATATTCCAT-3′ and 5′-
TCCTCCCACCTCGGCCTCCCAAACTGCTGGGATTACAGATGTGAGCCACTGCACC
CAGCCGGTAACAGAGTATTTTAAAATAACATCATATTCCATTATTGTTCAGGATCTT
GGAACATTGAGTGATATA-3′. Ribonucleoprotein (RNP) complexes were assembled by incubating 1.5 μg of recombinant Cas9 protein with 360 ng of each gRNA at room temperature for 10 minutes. The resulting RNPs were combined with the ssODN repair templates and electroporated into $1\times10^5$ iPSCs using the Neon Transfection System (Thermo Fisher Scientific). Transfected cells were subsequently plated in Geltrex-coated 6-well plates and cultured in StemFlex™ medium, supplemented with CloneR™ (Stemcell Technologies). Genomic DNA was extracted three days after transfection to screen for the T>C edit. PCR

amplification was performed using primers 5′-CTAACTGCAGCCTCTGTCTC-3′ and 5′-CATCCCTTCACTCTGATCCATTA-3′, followed by Sanger sequencing to confirm the presence of the desired base substitution. Clonal lines were derived by single-cell sorting and expanded for further analysis. Sanger sequencing was used to confirm the genotype of each clone. Homozygous (C/C) clones BIHi005-A-6 (005-A/1 herein) and BIHi005-A-7 (005-A/2 herein) from parental line BIHi005-A as well as MDCi053-A-58 (053-A/1 herein) and MDCi053-A-60 (005-A/2 herein) from parental line MDCi053-A-49 were chosen for further analysis. Prior, clones underwent quality control and were routinely tested to be free of mycoplasma contamination.

**_SORL1_ and _APOE_ genotyping**

Genomic DNA was isolated using Genomic DNA purification Kit (Promega #A1120) according to the manufacturer's protocol and subjected to quantitative PCR using the TaqMan SNP Genotyping Assay and TaqMan Genotyping Master Mix (Applied Biosystems). Genotypes were determined based on SNP rs429358 that defines ε3 and ε4 alleles of human *APOE* (Assay ID C_3084793_20, Applied Biosystems) or rs11218343 that defines T and C alleles of human *SORL1* (Assay ID C_31696474_10, Applied Biosystems). FAM and VIC reporter dyes were used for allele discrimination.

**Differentiation of human iPSCs into induced neurons**

Human iPSC lines were differentiated into induced neurons (iNs) using lentiviral vectors encoding *NGN2*, *rtTA*, and *EGFP* (2). Cells were cultured in F12-N2 medium supplemented with 1X N2 (Thermo Fisher #17502-048), 1X NEAA (Thermo Fisher #11140-035), 10 ng/ml human BDNF (R&D Systems #248-BD-025), 10 ng/ml human NT3 (R&D Systems #267-N3-025), 0.2 μg/ml mouse laminin (Thermo Fisher #23017-015) and 2 μg/ml doxycycline

(Sigma-Aldrich #D9891). Doxycycline was retained in the culture medium throughout the experiment to sustain expression of *NGN2* via the doxycycline-inducible system. Puromycin (8 µg/ml) was added to the medium to select for cells transduced with lentiviral vector expressing *NGN2* (2). The medium was gradually exchanged to NB-B27 medium (Thermo Fisher #21103-049 Neurobasal medium), containing 1X B27 (Thermo Fisher #17504-044), 1X GlutaMAX (Thermo Fisher 35050-061), 10 ng/ml human BDNF (R&D Systems), 10 ng/ml human NT3 (R&D Systems), and Ara-C (Sigma-Aldrich #C1768). Induced neurons were maintained in culture for 14 days before analysis.

**Differentiation of human iPSCs into induced astrocytes**

Induced astrocytes (iAs) were generated from human iPSCs using lentiviral vectors encoding *NFIB*, *SOX9*, *rtTA*, *mCherry* (3). Cells were cultured in expansion medium (Thermo Fisher # 31330-038 DMEM/F12) containing 1X N2 (Thermo Fisher), 1X GlutaMAX (Thermo Fisher), 10% FBS (Thermo Fisher #10082-147), and 2,5 µg/ml doxycycline. Doxycycline was kept in the medium until the end of the experiment. Puromycin (0.25 µg/ml) and hygromycin (200 µg/ml) were added to the culture medium to select for cells transduced with lentiviral vectors expressing *SOX9* and *NFIB* (3). The medium was gradually exchanged to FGF medium composed of Neurobasal medium (Thermo Fisher), containing 2% B27 (Thermo Fisher), 1X NEAA (Thermo Fisher), 1X GlutaMAX (Thermo Fisher), 1% FBS (Thermo Fisher), 8 ng/ml human FGF (Peprotech #100-18B), 5 ng/ml human CNTF (Peprotech #450-13), and 10 ng/ml human BMP4 (Peprotech #120-05ET). Cells were dissociated with Accutase and cultured in FGF medium. Thereafter, half of the medium was replaced every other day by maturation medium composed of a 1:1 mixture of DMEM/F12 (Thermo Fisher) and Neurobasal medium (Thermo Fisher), containing, 1X N2 (Thermo Fisher), 1X GlutamMAX (Thermo Fisher), 1% sodium pyruvate (Thermo fisher ##11360-

039), 5 µg/ml N-acetyl cysteine (Sigma-Aldrich #A8199), 500 µg/ml dbcAMP (Sigma-Aldrich #D0627), 5 ng/ml EGF-like growth factor (Sigma-Aldrich #E4643), 10 ng/ml CNTF (Peprotech), and 10 ng/ml BMP4 (Peprotech). Induced As were maintained for 28 days before analysis.

**Differentiation of human iPSCs into induced microglia**

Induced microglia (iMg) were generated from human iPSCs as described previously (4). In brief, iPSCs were differentiated into hematopoietic progenitor cells (HPs) using the STEMdiff™ Hematopoietic Kit (Stem Cell Technologies #05310). HPs (day 11) were harvested and resuspended in differentiation medium (Thermo Fisher #11039-021 DMEM/F-12) comprised of 2X Insulin-Transferrin-Selenite (Thermo Fisher #51300-044), 2X B27 (Thermo Fisher), 0.5X N2 (Thermo Fisher), 1X GlutaMAX (Thermo Fisher), 1X NEAA (Thermo Fisher), 400 µM monothioglycerol (Sigma-Adrich #M1753), 5 µg/ml insulin (Sigma-Aldrich #I2643), 100 ng/ml IL-34 (PeproTech #200-34), 50 ng/ml TGFβ1 (1PeproTech #00-21C), and 25 ng/ml M-CSF (PeproTech #300-25). From days 13 to 21, fresh medium was added to replenish the volume every other day. At day 23, medium was centrifuged, the supernatant was removed and the cell pellets were resuspended in fresh differentiation medium. From days 25 to 33, fresh differentiation medium was added to replenish volume every second day. At day 35, iMg were transferred to maturation medium containing 100 ng/ml CD200 (Elabscience #E-PKSH032840), and 100 ng/ml CX3CL1 (PeproTech #300-31) and supplemented with fresh medium every other day until day 38. Between days 38-39 of further culture, iMg were used for functional studies.

**Microglia stimulations**

For pro-inflammatory stimulation, iMg were harvested, replated in microglia differentiation medium, and stimulated with 2,5 µg/ml, 5 µg/ml or 10 µg/ml Poly(I:C) (Sigma-Aldrich

#P9582) for 24h. Cultures treated with PBS in differentiation medium served as unstimulated solvent control. Cytokine levels in medium samples were determined using ELISA kits for human TNFα, IL-1β, IL-6, and RANTES (K15231N-1, K15067L-1, U-Plex, Mesoscale) as well as human IFN-β (151ADRS-1, S-Plex, Mesoscale). Total RNA was isolated from stimulated iMg (RNeasy Plus Micro Kit, Qiagen #74034), and used for transcription analysis.

**Immunocytochemistry**

Cells were fixed with 4% PFA in PBS and blocked with 5% normal donkey serum, 0.25% Triton X-100 in PBS at room temperature for 30 min, followed by incubation with primary antibodies in blocking buffer at 4°C overnight: anti-IBA1 (diluted 1:100, Abcam #ab5076), and anti-P2RY12 (diluted 1:50, Sigma-Aldrich #HPA013796). The corresponding secondary antibodies (diluted 1:1000) in blocking buffer were used (for 2 hours at room temperature). Nuclei were counterstained with DAPI (1:500, Anaspec #AS-83211). Images were acquired using a Zeiss LSM780 confocal microscope at 20x magnification, and analyzed using FIJI software.

**Protein expression analysis**

Cells were lysed in RIPA buffer (50 mM Tris-HCl pH 7.4, 150 mM NaCl, 0.5% sodium deoxycholate, 0.1% SDS 1% NP-40) containing 1X complete protease inhibitor cocktail (Roche #11697498001). An equal amounts of proteins from the various experimental conditions were subjected to standard SDS-PAGE, followed by transfer to PVDF membranes (Bio-Rad #162-0184). Membranes were blocked with 5% nonfat milk in TBS (with 0.5% Tween-20) and incubated with the following primary antibodies in TBST: anti-SORLA (diluted 1:1000, BD Biosciences #611861) or anti-GAPDH (diluted 1:3000, Sigma-Aldrich #MAB374). Corresponding secondary antibodies conjugated to horseradish peroxidase were used (diluted 1:4000) and membranes were developed on iBright CL1500 Imaging System (Invitrogen). Densitometric analysis was performed using the FIJI software. For each sample, the intensity of SORLA immunoreactive band was normalized to the GAPDH signal.

**Transcription analysis**

Total RNA was extracted from cell cultures using RNeasy Plus Mini/Plus Micro Kit (Qiagen #74134, 74034), treated with RNase-Free DNase Set (Qiagen #79254), and cDNA synthesis was performed with High Capacity RNAto-cDNA Kit (Thermo Fisher #4374967). Quantitative PCR was conducted on a QuantStudio 7 Flex Real-Time PCR System (Applied Biosystems) using TaqMan gene expression assays for *SORL1* (Hs00983770), *OCT4* (Hs00999632_g1), *SOX2* (Hs01053049_s1), *NANOG* (Hs02387400_g1), *TUBB3* (Hs00964963_g1), *MAP2* (Hs00258900_m1), *IBA1* (Hs00610419), *P2RY12* (Hs01881698), *GFAP* (Hs00909233_m1), *IL-1β* (Hs01555410_m1), *IL-6* ( Hs00174131_m1), *IFNβ* (Hs01077958_s1) *RANTES* (Hs00982282_m1) *TNFα* (Hs00174128_m1), *GAPDH* (Hs02758991_g1), *TBP* (Hs00427620_m1). The fold change in transcript levels was determined using the cycle threshold (CT) comparative method ($2^{-ddCT}$). Data were normalized to the reference genes *GAPDH* (for iNs and iAs) or *GAPDH* and *TBP* (for iMg). To assess genotype-dependent differences in transcript levels, ΔCt values for each T/T, T/C and C/C cell line were normalized to the mean of ΔCt values of all T/T cell lines in the data set (set to relative level 1).

SUPPLEMENTARY FIGURES AND  LEGENDS

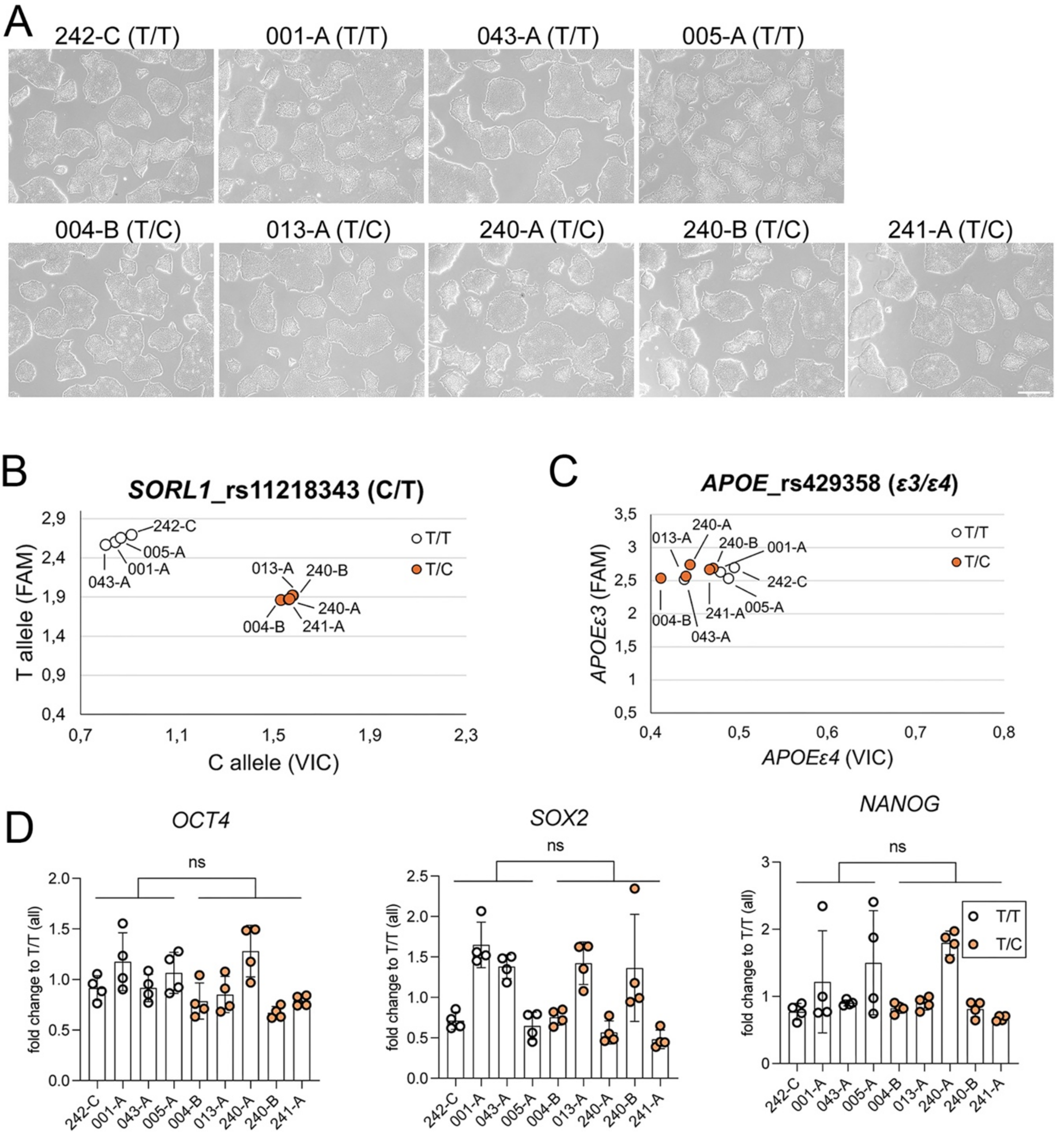


**Supplementary figure 1: Characterization of the donor iPSC lines**

(**A**) Representative bright field images of human donor iPSC lines carrying the indicated T/T or T/C alleles of rs11218343. Scale bar: 500 µm. (**B)** SNP genotyping of donor iPSC lines for rs11218343. Scatter plots represent allele discrimination for T/T (white dots) and T/C (orange dots) genotypes. (**C**) SNP genotyping of donor iPSC lines for *APOEε3* and *APOEε4*. All donor iPSC lines are *APOEε3/APOEε3*. (**D**) Quantitative RT-PCR analysis of transcript levels for pluripotency markers *OCT4*, *SOX2*, and *NANOG* in the indicated donor iPSC lines. Data are shown as mean ± SD of 4 biological replicates (from 4 independent differentiations) for each cell line. Statistical significance of data was determined using nested *t* test comparing genotypes. ns, not significant.

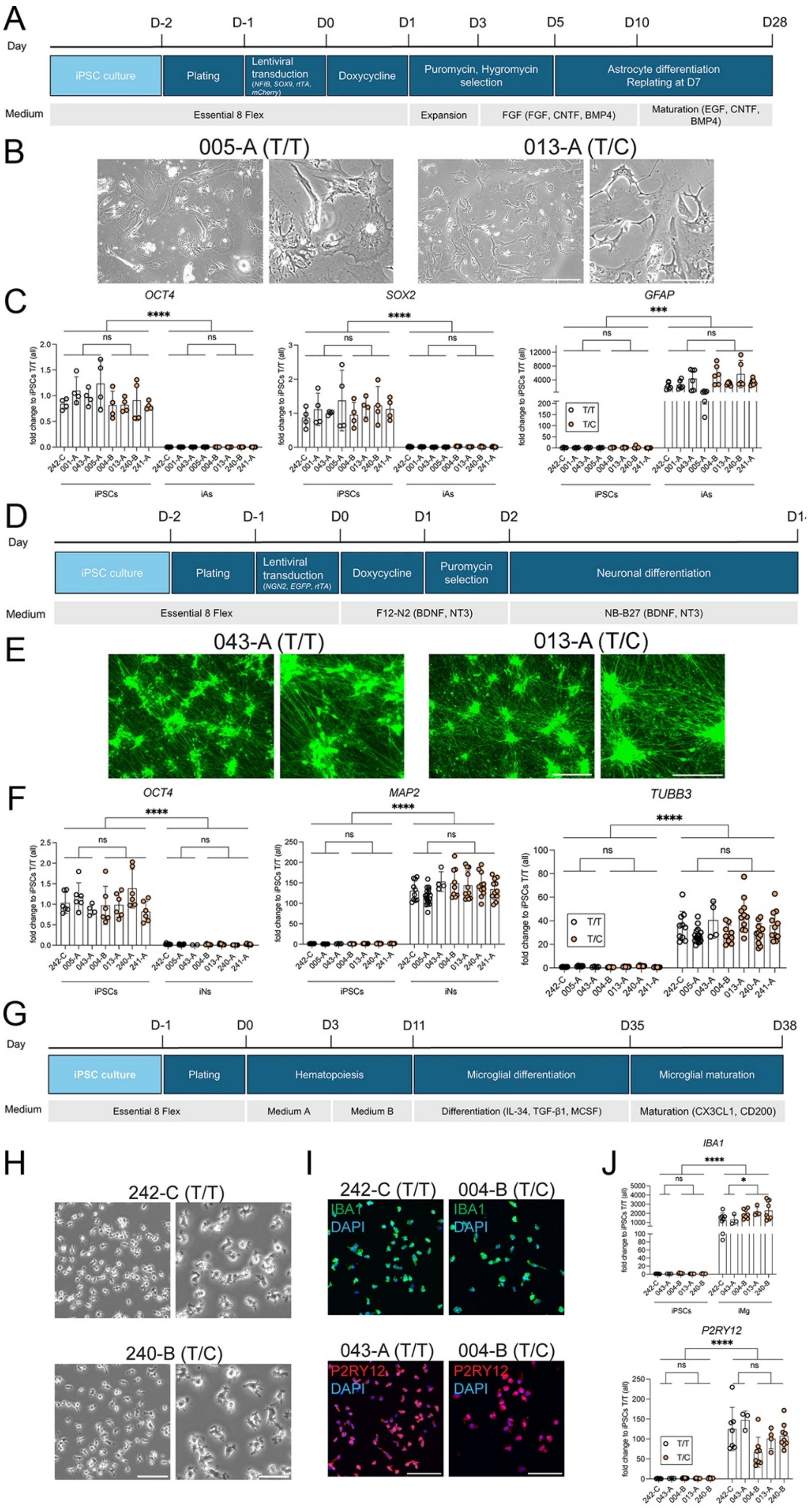


**Supplementary figure 2: Differentiation of donor iPSC lines into human astrocytes, neurons, or microglia**

(**A**) Protocol for differentiation of human donor iPSC lines into astrocytes (iAs). (**B**) Exemplary bright field images of iAs cultures of one T/T and one T/C genotype at day 28 of differentiation. Scale bars: 300 µm and 150 µm (zoom-in). (**C**) Quantitative RT-PCR analysis of transcript levels of pluripotency markers *OCT4* and *SOX2,* as well as astrocyte marker *GFAP* in the indicated cell lines as iPSC and when differentiated to iAs. Data are shown as mean ± SD of 4 biological replicates from 4 independent differentiations (for iPSC) or 5-7 biological replicates from 3 independent differentiations (for iAs) for each cell line. Statistical significance of data was determined using nested *t* test comparing transcript levels in iPSC and iAs cell types or between genotypes within each cell type. ns, not significant; ***, $p < 0.001$; ****, $p < 0.0001$. (**D**) Protocol for differentiation of iPSC lines into induced human neurons (iNs). (**E**) Immunofluorescence images of human iNs derived from two exemplary donor iPSC lines of T/T or T/C genotype (day 14 of differentiation). Neurons were visualized by native fluorescence of GFP, encoded by an *EGFP* expression construct included in the neuronal differentiation protocol (see methods for details). Scale bars: 300 µm and 150 µm (zoom-in). (**F**) Quantitative RT-PCR analysis of transcript levels for pluripotency marker *OCT4* as well as neuronal markers *MAP2* and *TUBB3* in the indicated cell lines as iPSC and when differentiated to iNs. Data are the mean ± SD of of 4-6 biological replicates from 4-6 independent differentiations (for iPSC) or 4-18 biological replicates from 3-11 independent differentiations (for iNs) for each donor line. Statistical significance of data was tested as described in panel (C). (**G**) Protocol for differentiation of donor iPSC lines into induced human microglia (iMg). (**H**) Bright field images of human iMg cells derived from two exemplary iPSC lines with T/T or T/C genotype at day 38 of differentiation. Scale bars: 100 µm and 50 µm (zoom-in). (**I**) Immunostaining of two exemplary human iMg lines for microglia marker IBA1 (green) or P2RY12 (red). Nuclei were counterstained with DAPI (blue). Scale bars: 100 µm. (**J**) Quantitative RT-PCR analysis of transcript levels for *P2RY12* and *IBA1* in iMg lines of the indicated T/T and T/C genotypes. Data are the mean ± SD of 3-6 biological replicates from 3-6 independent differentiations (iPSC) or 3-9 biological replicates from 3-9 independent differentiations (iMg) for each iPSC line. Statistical significance of data was tested as described in panel (C).

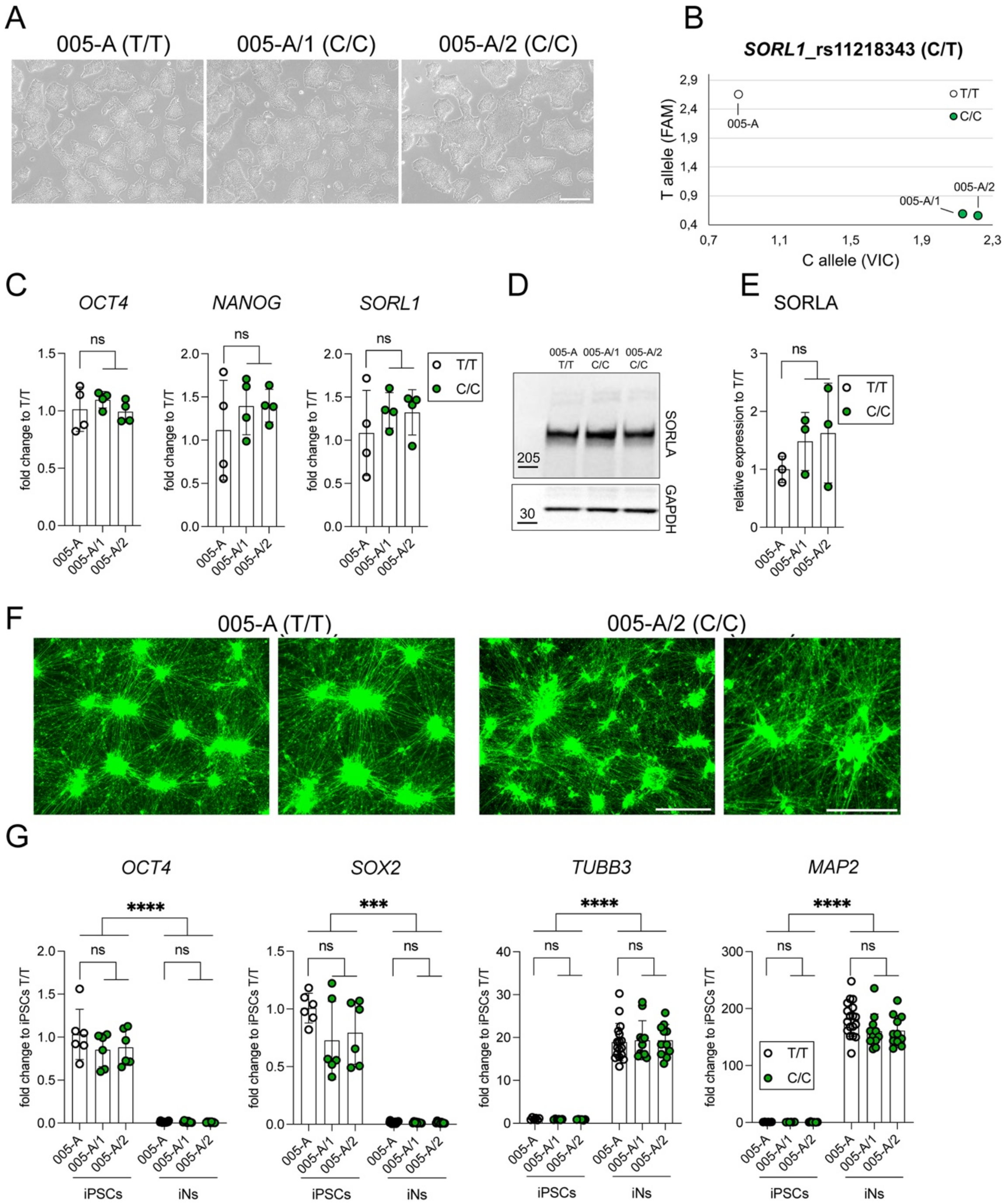


**Supplementary figure 3: Characterization of isogenic iPSC lines and induced human neurons, genome-edited for rs11218343**

(**A**) Representative bright field images of parental iPSC line 005-A (T/T) and two independent isogenic clones 005-A/1 and /2, genome-edited to carry the C/C genotype. Scale bar: 500 µm. (**B**) SNP genotyping of the indicated iPSC lines for rs11218343. Scatter plots represent allele discrimination for T/T (white dots) and C/C (green dots) genotypes. (**C**) Quantitative RT-PCR analysis of transcript levels for *OCT4*, *NANOG*, and *SORL1* in the indicated iPSC lines. Data are shown as mean ± SD of 4 biological replicates (from 4 independent differentiations) for each cell line. Statistical significance was determined using nested *t* test comparing genotype groups. ns, not significant. (**D, E**) Levels of SORLA in the indicated iPSC lines were determined by Western blotting of cell lysates (D) and densitometric scanning of replicate blots (E). Relative levels of SORLA were normalized to the GAPDH loading control in each sample. Data are the mean ± SD of 3 biological replicates (from 3 independent differentiations) for each cell line. Statistical significance was determined using nested *t* test comparing genotypes. (**F**) Immunofluorescence images of human iNs derived from isogenic cell lines 005-A (T/T) and 005A/2 (C/C) at day 14 of differentiation. ). Neurons were visualized by native fluorescence of GFP, encoded by an *EGFP* expression construct included in the neuronal differentiation protocol (see methods for details). Scale bars: 300 μm and 150 μm (zoom-in). (**G**) Quantitative RT-PCR analysis of transcript levels for pluripotency markers *OCT4* and *SOX2*, as well as neuronal markers *TUBB3* and *MAP2* in the indicated iPSC lines and iNs derived thereof. Data are the mean ± SD of 6 biological replicates from 6 independent differentiations (for iPSC) or 11-18 biological replicates from 8-11 independent differentiations (for iNs) for each cell line. Significance of data was determined using nested *t* test when comparing transcript levels in iPSC and iNs cell types or between genotypes within each cell type. ***, $p < 0.001$; ****, $p < 0.0001$.

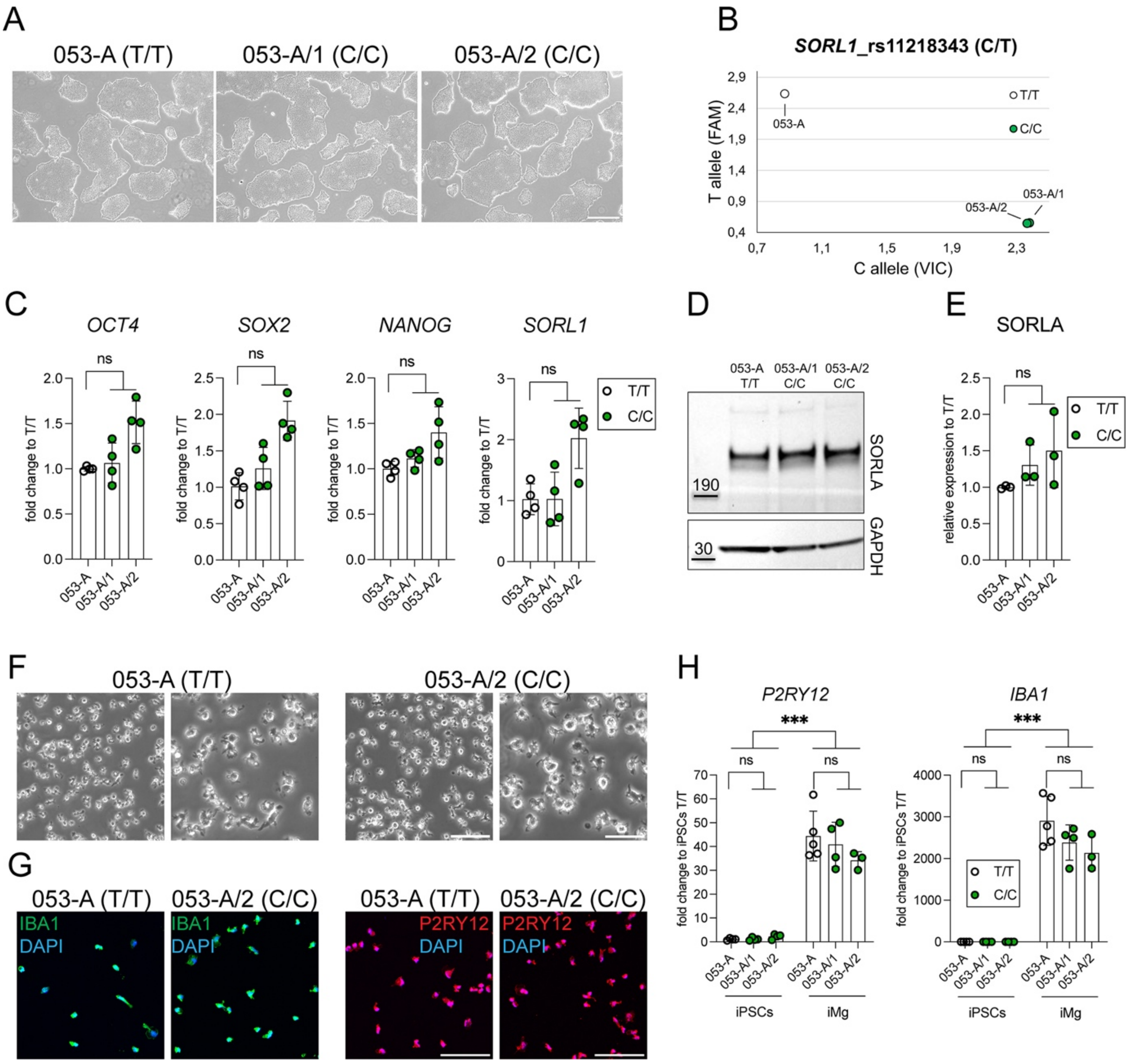


**Supplementary figure 4: Characterization of isogenic iPSC lines and induced human microglia, genome-edited for rs11218343**

(**A**) Representative bright field images of parental iPSC line 053-A (T/T) and two independent isogenic cell clones 053-A/1 and /2, genome-edited to carry the C/C genotype. Scale bar: 500 µm. (**B**) SNP genotyping of the indicated iPSC lines for rs11218343. Scatter plots represent allele discrimination for T/T genotype (white dot) and C/C genotype (green dots) of rs11218343. (**C**) Quantitative RT-PCR analysis of transcript levels for *OCT4*, *SOX2*, *NANOG*, and *SORL1* in the indicated iPSC lines. Data are shown as mean ± SD of 4 biological replicates (from 4 independent differentiations) for each cell line. Statistical significance of data was determined using nested *t* test comparing genotypes. ns, not significant. (**D, E**) Levels of SORLA in the indicated iPSC lines were determined by Western blotting of cell lysates (D) and densitometric scanning of replicate blots (E). Relative levels of SORLA were normalized to the GAPDH loading control in each sample. Data are the mean ± SD of 3 biological replicates (from 3 independent differentiations) for each cell line. Statistical significance of data was determined using nested *t* test comparing genotypes. (**F**) Representative bright field images of isogenic iMg cultures of the indicated T/T and the C/C genotypes at day 38 of differentiation. Scale bars: 100 µm and 50 µm (zoom-in). (**G**) Immunostaining of two exemplary human iMg lines for microglia marker IBA1 (green) or P2RY12 (red). Nuclei were counterstained with DAPI (blue). Scale bars: 100 µm. (**H**) Quantitative RT-PCR analysis of transcript levels for microglia marker genes *P2RY12* and *IBA1* in the indicated iPSC lines and in iMg derived thereof. Data are the mean ± SD of 4 biological replicates from 4 independent differentiations (for iPSC) or 3-5 biological replicates from 3-5 independent replicates (for iMg). Significance of data was determined using nested *t* test comparing transcript levels in iPSC and iMg cell types or between genotypes within each cell type. ***, $p < 0.001$.
